# SMAC mimetics induce human macrophages to phagocytose live cancer cells

**DOI:** 10.1101/2024.11.25.625306

**Authors:** Samantha Y. Liu, Max Hulsman, Philipp Leyendecker, Eugena Chang, Katherine A. Donovan, Fabian Strobel, James Dougan, Eric S. Fischer, Michael Dougan, Stephanie K. Dougan, Li Qiang

## Abstract

Macrophages engulf apoptotic bodies and cellular debris as part of homeostasis, but they can also phagocytose live cells such as aged red blood cells. Pharmacologic reprogramming with the SMAC mimetic LCL161 in combination with T cell-derived cytokines can induce macrophages to phagocytose live cancer cells in mouse models. Here we extend these findings to encompass a wide range of monovalent and bivalent SMAC mimetic compounds, demonstrating that live cell phagocytosis is a class effect of these agents. We demonstrate robust phagocytosis of live pancreatic and breast cancer cells by primary human macrophages across a range of healthy donors. Unlike mouse macrophages where combination of SMAC mimetics with lymphotoxin enhanced phagocytosis, human macrophages were more efficiently polarized to phagocytose live cells by the combination of SMAC mimetics and IFNγ. We profiled phagocytic macrophages by transcriptional and proteomic methodologies, uncovering a positive feedback loop of autocrine TNFα production.

---

Macrophage clearance of apoptotic cells, also known as efferocytosis, plays a critical role in eliminating damaged cells to maintain homeostasis. Efferocytotic macrophages also promote tissue repair through secretion of growth factors, suppression of effector T cell responses through IL-10 and TGFβ, and downregulation of inflammatory cytokines TNFα and IL1β ^1^. Macrophages distinguish apoptotic cells from live cells through scavenger receptors, including Tim1, Tim4, CD36 and CD300f, which can bind to phosphatidylserine on the outer layer of the apoptotic cells^1^. Bridging receptors like Mertk bind to Gas6 and Protein S that engage phosphatidylserine on dying cells ^2,3^. Macrophages can also secrete various factors to recognize dying cells and enhance efferocytosis, including Mfge8 that binds to lipids and calreticulin that binds to asialoglycoproteins on target dying cells and tags them for recognition by C1q and Lrp1 ^4–7^.

Live cells, on the other hand, are generally protected from being phagocytosed by “don’t eat me” signals. CD47/SIRPα is the best characterized pair in which CD47 expressed by healthy cells binds to the ITIM-domain containing receptor SIRPα on macrophages to prevent phagocytosis ^8–10^. Several other negative regulators of phagocytosis have been discovered including LILBR1/MHC class I and Siglec10/CD24 for human cells ^11,12^. Loss of CD47, particularly on aged erythrocytes precedes their clearance from circulation by red pulp macrophages in the spleen. CD47 expression is heterogeneous on epithelial cells and is upregulated on cancer cells, suggesting that prevention of phagocytosis may be a means of evading macrophage-mediated immune surveillance.

Regulation of phagocytosis of microbes is relatively well-understood. MBL, Dectin-1, and C1q all trigger particle engulfment. IgG antibodies bound to the surface of target cells can engage activating Fc receptors on macrophages and strongly induce phagocytosis ^13,14^. Pharmacologic antibodies against tumor cell surface antigens are effective therapeutics, and opsonization of cancer cells for engulfment by macrophages is likely part of the mechanism of action of these drugs. Complement deposition on target cells binds to CR3 (CD11b/CD18) on macrophages and induces phagocytosis via Syk and rearrangement of the actin cytoskeleton ^15^. Most healthy cells express complement decay factors but complement deposition on neurons and trogocytosis by macrophages is also critical for synaptic pruning during development of the CNS ^16^.

Endogenous positive regulators of cancer cell phagocytosis are still largely unknown. Live cancer cells have higher levels of surface exposed phosphatidylserine on their plasma membranes, even when not undergoing apoptosis, which may serve as a phagocytic target ^3,17^. Pro-phagocytic secreted calreticulin may also preferentially stick to the surface of tumor cells ^4–6^. Understanding whether and how macrophages can phagocytose live cancer cells while sparing surrounding healthy tissue is important to understanding how we could manipulate these pathways for therapeutic benefit.

The cellular inhibitor of apoptosis proteins (cIAPs) regulate NF-κB signaling and can be antagonized pharmacologically by peptide mimetics of SMAC, the natural binding partner of the IAP family ^18–21^. Several SMAC mimetics have been developed including monovalent and bivalent engagers as well as newer non-peptide versions ^22^. Although these agents were originally designed amplify autocrine TNFα production and extrinsic apoptosis pathways in cancer cells, their efficacy in syngeneic mouse cancer models appears to be primarily through induction of anti-tumor immunity ^23^. Degradation of cIAP1 by SMAC mimetics induces non-canonical NF-kB signaling and mimics co-stimulation through TNF receptor superfamily members such as GITR and 4-1BB to control cytokine output from activated T cells ^24–30^. SMAC mimetics also act directly on myeloid cells. The monovalent SMAC mimetic LCL161 was reported to induce phagocytosis of multiple myeloma cells despite genetic loss of *BIRC2* (the gene encoding cIAP1) from the cancer cells, thereby implicating macrophages as the cellular target of LCL161’s anti-cancer activity ^31^. In mouse models of pancreatic cancer, we identified that macrophages can engulf live tumor cells upon activation by LCL161 along with T cell–produced cytokines ^30,32^. In wild-type mice, but not mice incapable of antigen-specific T cell responses, cIAP1/2 antagonism reduces tumor burden by increasing phagocytosis of live tumor cells. This efficacy could be augmented by combination with CD47 blockade, suggesting that LCL161 activates tumoricidal macrophages independently of CD47/SIRPα, potentially via activation of pro-phagocytic pathways ^32^.

Thus far, most of the experiments showing SMAC mimetic induction of live cell phagocytosis have been conducted using LCL161 in mice or in mouse bone marrow-derived macrophages ^31,33,34^. Here we used an in vitro phagocytosis assay to evaluate a panel of structurally divergent SMAC mimetic compounds in primary monocyte-derived macrophages from a range of healthy human donors. Phagocytosis could be induced by all SMAC mimetics tested and was enhanced by co-culturing with IFNγ. The transcriptional signature of phagocytic macrophages was dominated by a subset of NF-κB target genes and sustained by autocrine TNFα production.

## Materials and Methods

### Ethics statement

These studies were conducted in accordance with the Declaration of Helsinki and the Belmont Report. Anonymous healthy donor leukopacks were obtained from the Kraft Family Blood Donor Center at Dana-Farber Cancer Institute and Brigham and Women’s Hospital, protocol T0363.

### Cell culture

Both MDA-MB-231 and PANC-1 cell lines were cultured in RPMI 1640 Medium (Thermo Fisher, Cat# 11875135) supplemented with 10% heat-inactivated fetal bovine serum, Penicillin-Streptomycin 100 U/mL (Thermo Fisher, Cat# 15140122), MEM Non-Essential Amino Acids Solution (Thermo Fisher, Cat #11140050), GlutaMAX Supplement (Thermo Fisher, Cat# 35050061) and Sodium pyruvate (100 mM) (Thermo Fisher, Cat# 11360070), i.e. RMPI complete. Cell lines were cultured at 37°C in a humidified incubator containing 5% carbon dioxide (CO2).

### Peripheral blood mononuclear cell (PBMC) isolation and macrophage differentiation

PBMCs were derived from healthy donor leukopacks using density gradient centrifugation with Ficoll-Paque PLUS Media (Fisher Scientific, Cat# 45001749) in SepMate™ PBMC Isolation Tubes (Stemcell Technologies, Cat# 85450). Prior to centrifugation the whole blood was diluted with an equal volume of PBS, pH 7.4 (Thermo Fisher, Cat# 10010049). Upon centrifugation the buffy coat containing PBMCs was collected and red blood cell lysis was performed using ACK lysis buffer (Thermo Fisher, Cat# A1049201). Subsequently, PBMCs were resuspended in RPMI complete supplemented with recombinant human M-CSF (hM-CSF) 50 ng/mL (PeproTech, Cat# 300-25) and plated on 150 x 15 mm petri dishes (Fisher Scientific, Cat# 08-757-148) at a concentration of 1–2 million cells per mL. In order for proper differentiation of PBMCs into macrophages, cells were fed on day 2 and 4 of culture by adding the same volume of RPMI complete with 50 ng/mL of recombinant human M-CSF.

### Phagocytosis assay

We performed in vitro live cell phagocytosis assays as described ^33^. Upon 5-6 days of differentiation, macrophages were collected from the petri dishes using 15 ml of 10% Trypsin-EDTA (0.25%), phenol red (Gibco, Cat# 25200056) in PBS. Subsequently, macrophages were again resuspended in hM-CSF supplemented RMPI complete and reseeded into 12-well tissue culture-treated plates (Fisher Scientific, Cat# 07-200-82) by adding 1 mL into each well of the 12-well plates at 100,000 cells/mL. Macrophages were incubated at 37°C in a humidified incubator containing 5% CO2 and used for phagocytosis assessment at days 7-12 of culture. Prior to the start of the phagocytosis assays, macrophages were pre-treated for 24 hours with various SMAC mimetics including ASTX-660 (MedChemExpress, Cat# HY-109565), Birinapant (MedChemExpress, Cat# HY-16591), CUDC-427 (MedChemExpress, Cat# HY-15835), GDC-0152 (MedChemExpress, Cat# HY-13638), LCL161 (MedChemExpress, Cat# HY-15518) and Xevinapant (MedChemExpress, Cat# HY-15454), or DMSO (Sigma-Aldrich, Cat# D5879) as a vehicle control. Additionally, recombinant human lymphotoxin α1/ β2 (rhLTα1/β2) (R&D Systems, Cat# 8884-LY) or recombinant human IFNγ (PeproTech Cat# 300-02) was added during pre-treatment and co-culture. In some cases, cultures were also treated with anti-TNFα (InvivoGen, adalimumab biosimilar #htnfa-mab4). The next day, tumor cells were collected and labelled with CellTrace Violet (CTV) (Fisher Scientific, Cat# C34557) according to the manufacturer’s instructions. Briefly, cells were incubated with 5 μM CTV for 20 minutes at 37°C, protected from light. After incubation, the cells were washed three times with PBS to remove excess dye. Labelled tumor cells were then resuspended in human M-CSF supplemented RPMI complete at a concentration of 100,000 cells/mL. Subsequently, the treatment-containing media was aspirated from the macrophages, and tumor cells were added by pipetting 1 mL of the tumor cell suspension in each well. i.e. macrophage-to-tumor cell ratio of 1:1. Pre-treated macrophages were cocultured with tumor cells in a humidified incubator containing 5% CO2 for 18 hours. To collect both non-adherent and adherent cells, the supernatant from each well was first collected. Secondly, adherent cells were trypsinized and collected from each well. Both non-adherent and adherent cell fractions were combined in corresponding flow cytometry tubes containing HBSS, calcium, magnesium (Thermo Fisher, Cat# 14025126). Finally, flow cytometry tubes were centrifuged, and cell pellets were stained for flow cytometric phagocytic assessment.

### Flow cytometry

Each sample was stained in 100 μL FACS buffer, constituted of PBS, heat-inactivated fetal bovine serum 2% EDTA (Fisher Scientific, Cat# 15575020) 2 mM, and 1 μL of PE/Dazzle™ 594 anti-human CD45 Antibody (Biolegend, Cat# 304052) or PE-CF594 anti-human CD45 Antibody (BD Horizon, Cat# 562279). Samples were stained for 20 min at 4°C. Following staining, 200 μL of 1% formalin (Millipore Sigma, Cat# HT501128) was added to the tubes to fix the samples. Tubes were stored at 4°C and protected from light until time of analysis on a SP6800 Spectral Analyzer (Sony).

### Immunoblotting

Differentiated macrophages were reseeded in 6-well tissue culture-treated plates (Corning, Cat# 3516) at a density of 500,000 cells per well. The next day, cells were treated with 500 nM SMAC-mimetics or DMSO as a vehicle control for 24 hours. Media was then aspirated, and cells were washed twice with PBS. To obtain protein lysates, 50 μL of RIPA buffer (Abcam, Cat# ab156034) supplemented with cOmplete™, Mini, EDTA-free Protease Inhibitor Cocktail (Roche, Cat# 11836170001) and Phosphatase Inhibitor Cocktail (Cell Signaling Technology, Cat# 5870) was added to each well and plates were incubated on ice for 30 minutes. Lysates were collected using cell scrapers (Corning, Cat# 3011), centrifuged at 13,000 rpm for 30 minutes at 4°C, and the supernatants were stored at - 80°C until further analysis. Protein concentration of the lysates was determined using the Micro BCA™ Protein Assay Kit (Thermo Scientific, Cat# 23235). For sample preparation, 30 μg of protein was subsequently boiled for 5 minutes in reducing Laemmli SDS sample buffer (Thermo Scientific, Cat# J61337-AD), followed by rapid cooling on ice. Samples were then loaded onto 4-20% Tris-Glycine gels (BioRad, Cat# 4561094) and ran at 80-120V for SDS-PAGE electrophoresis. Following separation, proteins were transferred to a PVDF membrane (BioRad, Cat# 1620175) using the mixed MW program on a Trans-Blot Turbo Transfer System (BioRad, Cat# 1704150) for immunoblotting. Membranes were blocked with 5% BSA (Sigma-Aldrich, Cat# A7030) in 0.1% Tween-20 TBS (TBST) prior to overnight incubation at 4°C with primary antibodies. The primary antibodies used were c-IAP1 (D5G9) Rabbit mAb (Cell Signaling Technology, Cat# 7065S) and XIAP Rabbit Ab (Cell Signaling Technology, Cat# 2042S), each diluted 1:1000 in 3% BSA TBS-T. Following primary antibody incubation, membranes were washed three times with TBS-T and incubated with Anti-rabbit IgG, HRP-linked Antibody (Cell Signaling Technology, Cat# 7074) diluted 1:5000 in TBS-T. After three additional washes with TBS-T, protein bands were visualized using Western Lightning ECL Pro detection agent (Revvity, Cat# NEL120001EA) and capturing chemiluminescence on a ChemiDoc™ Imaging System (BioRad, Cat# 12003153).

### Transcriptomics

PBMCs were isolated from eight individual donors and differentiated into macrophages. For each donor, macrophages were treated for 24 hours with either DMSO, rhLTα1/β2, IFNγ, LCL161 (500nM), LCL161 + rhLTα1/β2, or LCL161 + IFNγ. After treatment, macrophages were harvested, and total RNA was extracted using the RNeasy Mini Kit (Qiagen, Cat# 74104) according to the manufacturer’s protocol. (company). Additionally, phagocytosis assays were performed on treated macrophages from all donors to ensure treatment response.

### Proteomics

Human macrophages were cultured under standard conditions and treated for 6 hours with our series of SMAC-mimetics at a concentration of 500nM or with DMSO as a vehicle control. Pomalidomide treatment was included as a control targeting the unrelated E3 ligase cereblon. Following treatment, macrophages were harvested and pelleted. Cell pellets were subjected to global quantitative proteomic analysis ^35–38^.

### Data analysis

Flow cytometry data were analyzed using FlowJo v10.8.1 (BD Biosciences) and further processed in Prism v9.3.0 (GraphPad Software). Statistical analyses for all assays were performed using ordinary one-way ANOVA. Additionally, for the cytokine synergy experiment, Dunnett’s multiple comparison test was applied. Protein band intensities from western blot images were quantified using ImageJ, while RNA-seq and proteomics data were processed and analyzed in RStudio.

### Data availability

Bulk transcriptional profiling is available at the NIH Gene Expression Omnibus repository under accession number GSE282835.

## Results

### SMAC mimetics induce phagocytosis of live tumor cells by human macrophages

IAP antagonists like SMAC-mimetics were originally developed as cancer therapeutics sensitizing tumor cells to TNFα-mediated apoptosis. We used a panel of monovalent (LCL161, CUDC-427, GDC-0152, xevinapant), bivalent (birinapant) and non-peptidomimetic (ASTX-660) SMAC mimetics that have advanced to clinical testing in humans ^39–45^. To determine whether these agents induce live cancer cell phagocytosis, we cultured monocyte-derived macrophages from healthy donors with 500nM of each SMAC mimetic for 24 hours prior to addition of fluorescently labeled MDA-MB-231 breast cancer cells. Phagocytosis was measured by flow cytometry as the percentage of macrophages staining positively for the tumor cell fluorophore (**Figure 1A** and **Supplemental Figure 1**). Engulfed tumor cell fluorescent signal ranged from bright to dim, consistent with macrophages digesting fluorescent cargoes in their phagolysosomes. We tested phagocytosis induction by our SMAC mimetic panel across 8 different healthy donors. Notably, responsiveness to SMAC-mimetic treatment varied among donors, but comparison to vehicle-treated macrophages for eight donors showed a significant increase in phagocytosis for all SMAC mimetics (**Figure 1B**).

**Figure 1.**
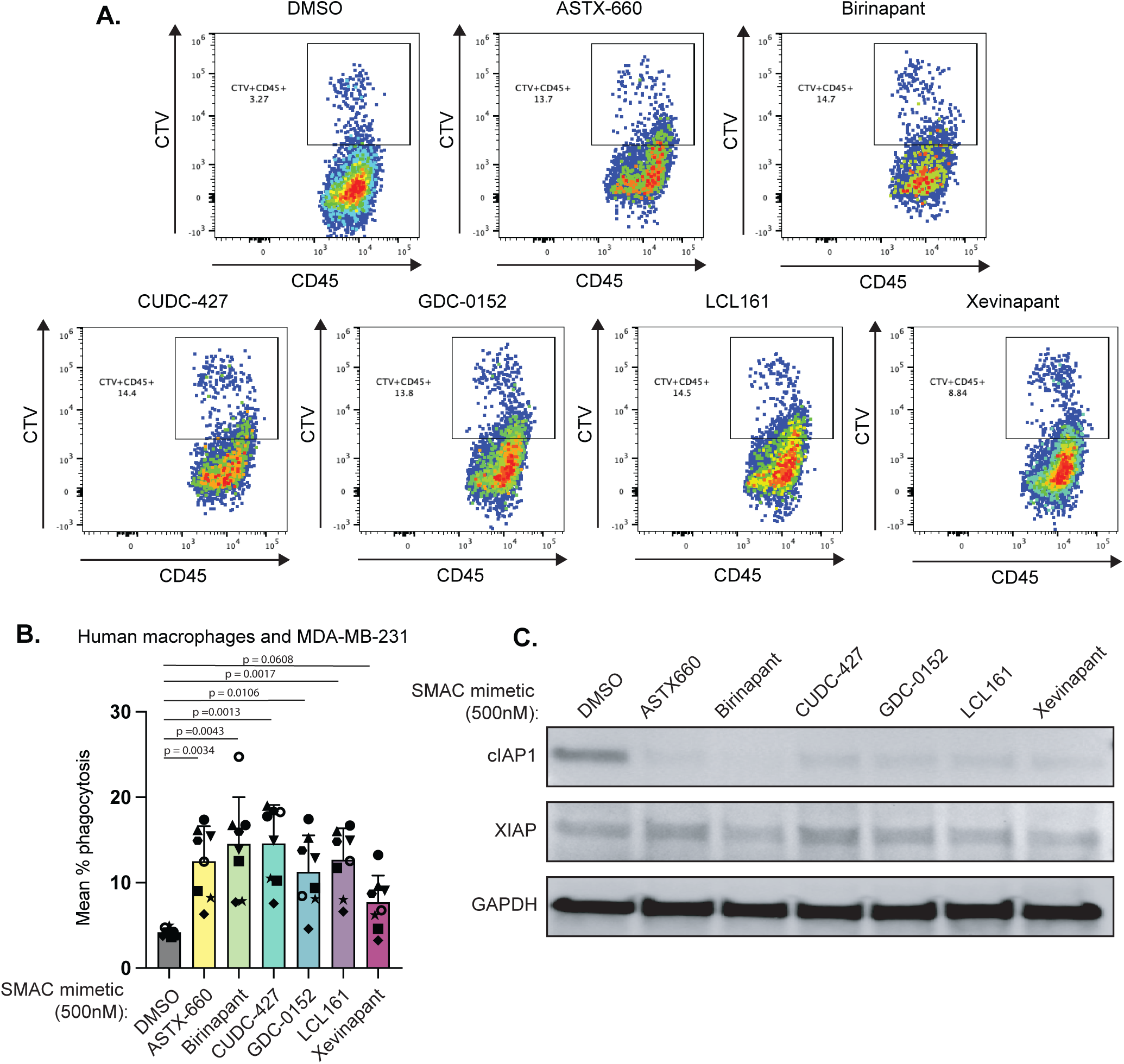
SMAC-mimetics induce phagocytosis in human macrophages. (A) Representative flow cytometry plots from a single donor showing the CTV⁺CD45⁺ phagocytosis population within the total CD45⁺ macrophage population. (B) Macrophages from n=8 healthy donors were treated with the 500nM of the indicated SMAC mimetics for 24 hours prior to co-culture with CellTrace Violet labeled MDA-MB-231 breast cancer cells. Phagocytosis was assessed by flow cytometry after 18 hours of co-culture. Results from each donor were normalized to the vehicle control from that donor. Error bars are SD. Statistical significance was determined using a pairwise one-way ANOVA with Sidak’s multiple comparison. (C) Immunoblot analysis of macrophage lysates treated with SMAC-mimetics (500 nM) for 24 hours. Blots were probed for cIAP1, XIAP, and GAPDH.

SMAC mimetics induce hyperactivation of the E3 ligase domain of cIAP1 and rapid auto-ubiquitination leading to degradation and stable loss of cIAP1 in treated cells ^46^. To confirm on-target activity of the SMAC mimetics at 500nM, we treated macrophages for 24 hours and analyzed cIAP1 and XIAP by immunoblot of protein lysates. Although cIAP1 protein was downregulated by all SMAC mimetics, XIAP was minimally affected (**Figure 1C**), consistent with prior reports showing that XIAP is more resistant to degradation by SMAC mimetics ^47^.

### Dose-dependent induction of phagocytosis in response to SMAC-mimetic treatment

To increase confidence that the enhanced phagocytic activity observed is a target-specific effect of SMAC mimetics and to identify the optimal concentration for maximal biological effect, we conducted a titration experiment in which human macrophages were treated for 18 hours with increasing concentrations of LCL161 and co-cultured with fluorescently labeled MDA-MB-231 tumor cells. Flow cytometry analysis revealed a dose-dependent increase in the percentage of phagocytosis+ macrophages, with the highest fold change in phagocytic activity at 500nM (**Figure 2A**). Consistent with these results, cIAP1 degradation rates increased with increasing concentration of LCL161, resulting in nearly undetectable levels of protein by immunoblot in lysates of macrophages treated with 500nM LCL161. More modest reductions were observed for XIAP1 (**Figure 2B**).

**Figure 2:**
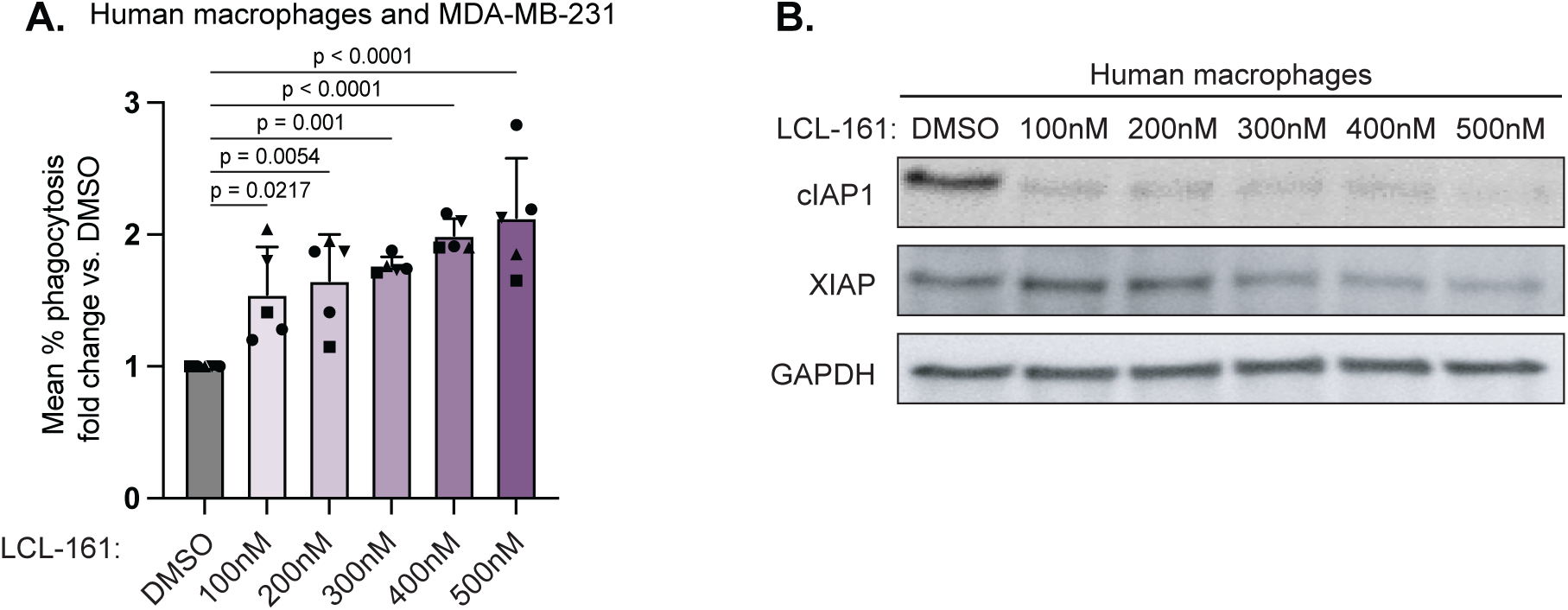
LCL161 induced phagocytosis is dose-dependent. (A) Macrophages from n=5 healthy donors were treated with the indicated concentrations of LCL161 for 24 hours prior to co-culture with CellTrace Violet labeled MDA-MB-231 breast cancer cells. Phagocytosis was assessed by flow cytometry after 18 hours of co-culture. Results from each donor were normalized to the vehicle control from that donor. Error bars are SD. Statistical significance was determined using an ordinary one-way ANOVA. (B) Immunoblot of protein lysates from macrophages treated for 24 hours with the indicated concentrations of LCL161.

### IFNγ enhances phagocytic of SMAC-mimetic treated macrophages

Building on our previous findings, which identified lymphotoxin as a critical cytokine for enhancing phagocytosis in cIAP1/2 antagonism therapy in murine models ^34^, we sought to explore whether T cell-produced cytokines enhanced human macrophage phagocytosis. We investigated the impact of lymphotoxin and IFNγ on macrophage-mediated phagocytosis using two different tumor cell lines: the breast cancer cell line MDA-MB-231 and the pancreatic cancer cell line PANC-1.

Our results indicate that IFNγ, rather than lymphotoxin, significantly enhances the phagocytic capacity of human macrophages, as shown in representative flow cytometry plots (**Figure 3A**). Notably, IFNγ treatment alone could modestly induce phagocytosis. In contrast to our previous observations in murine systems ^34^, lymphotoxin neither independently promoted phagocytosis nor enhanced the effects of LCL161 treatment. Importantly, the combination of LCL161 and IFNγ produced the most pronounced increase in phagocytic activity, as seen across multiple independent healthy donors and across two different cancer types (**Figure 3B-C**).

**Figure 3.**
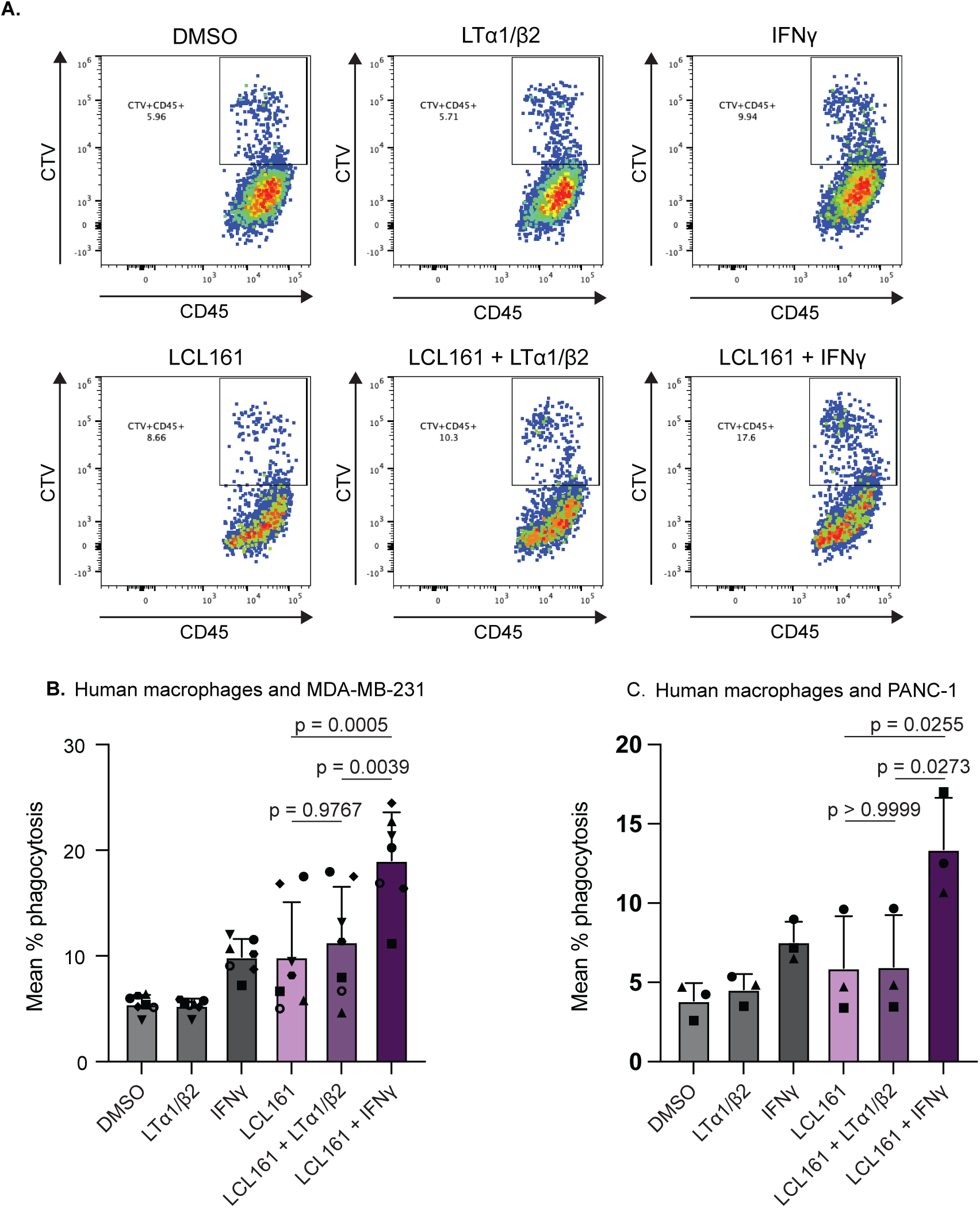
Phagocytic capacity of SMAC-mimetic treated macrophages is augmented by IFNγ. (A) Representative flow plots of human macrophages treated with LCL161 ± LTα1/β2 or IFNγ cocultured with CTV-labelled MDA-MB-231 tumor cells. Phagocytosis is shown as percentage of CTV+CD45+ macrophages within the total CD45+ population. (B) Quantified and analysed flow data from healthy donor macrophages (n=7 individual donors) co-cultured with MDA-MB-231 tumor cells. Error bars are SD. Statistical analysis was performed using an ordinary one-way ANOVA with Dunnett’s multiple comparison test. (C) Macrophages from n=3 healthy donors were treated as in B and co-cultured with PANC-1 tumor cells. Error bars are SD. Statistical analysis was performed using an ordinary one-way ANOVA with Dunnett’s multiple comparison test.

### Short-term proteomic effects are conserved across SMAC mimetics

SMAC mimetics activate the E3 ligase domain of cIAP1, thereby inducing rapid changes in protein abundance. We profiled human macrophages treated with our panel of SMAC mimetics for 6 hours to measure these short-term changes. As expected, protein levels of cIAP1 (*BIRC2*) decreased in all samples (**Figure 4**). Other proteins associated with the TNFR1/2 signaling complexes were not affected, suggesting that changes induced by altered NF-κB signaling require longer than 6 hours to occur, consistent with an 18-hour delay in acquisition of the phagocytosis phenotype. In addition to cIAP1/*BIRC2*, short term protein level decreases were observed for several secreted factors associated with macrophage activation including complement proteins, the neutrophil recruiting chemokine CXCL1, IL-1β, and ACOD1, the enzyme responsible for itaconate production. COX-2/*PTGS2*, was also decreased, suggesting a decrease in prostaglandins.

**Figure 4:**
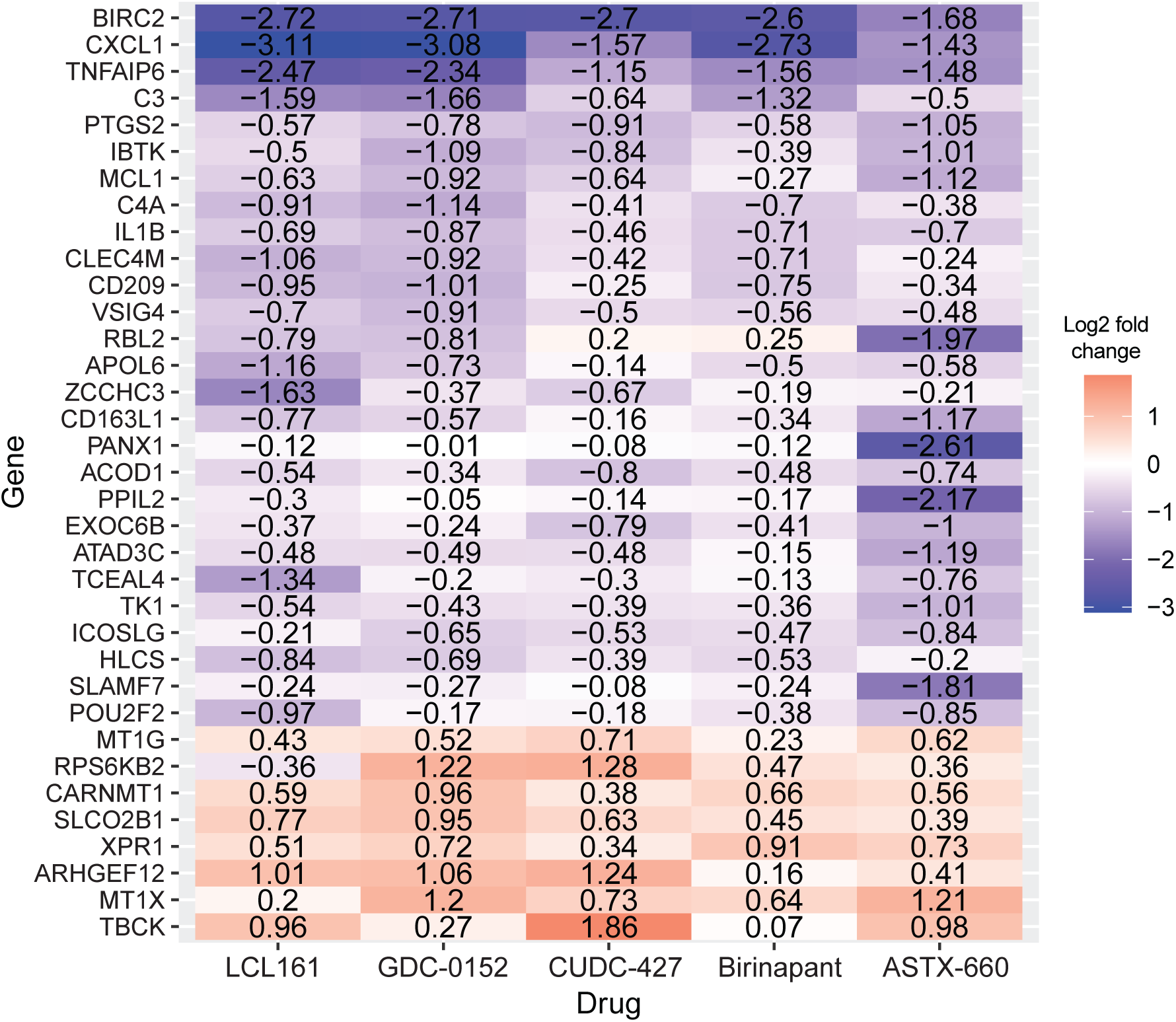
Short-term proteomic changes induced by SMAC mimetics. Healthy donor macrophages were treated for 6 hours with 500nM of the indicated SMAC mimetics, DMSO vehicle control, or pomalidomide, which targets the unrelated E3 ligase cereblon. Proteomic analysis was performed on n=6 technical replicates and reported as fold change compared to the DMSO control. Differentially expressed proteins were defined as having an absolute value of the average log2 fold change of the SMAC mimetics >0.5 and an absolute value of the average log2 fold change of pomalidomide <0.25.

### Transcriptional signature of phagocytosis is robustly conserved across human donors

Phagocytosis induction by SMAC mimetics requires at least 18 hours of culture, suggesting that transcriptional reprogramming underlies the phenotype. To define a transcriptional signature of phagocytosis, we cultured macrophages from 8 different healthy donors with vehicle, LCL161, IFNγ, or combination of LCL161 and IFNγ. RNA was collected at 24 hours and analyzed by bulk transcriptional profiling. Principle component analysis revealed clear separation between the two groups of samples treated with IFNγ versus those without, consistent with macrophages responding strongly to this cytokine (**Figure 5A**). The effect of LCL161 was more subtle, but comparison of samples from each donor shows an LCL-induced increase along the PCA2 axis (**Figure 5A**). Differential gene expression analysis for each treatment condition compared to vehicle control similarly shows that LCL161 alone induces a small number of NF-κB pathway genes (*MAP3K14*, *TRAF1, NFKB2, NFKBIE*) whereas treatment with IFNγ or LCL161/IFNγ induces a robust IFN response (*JAK2, CXCL9, CXCL10*) (**Figure 5B**).

**Figure 5:**
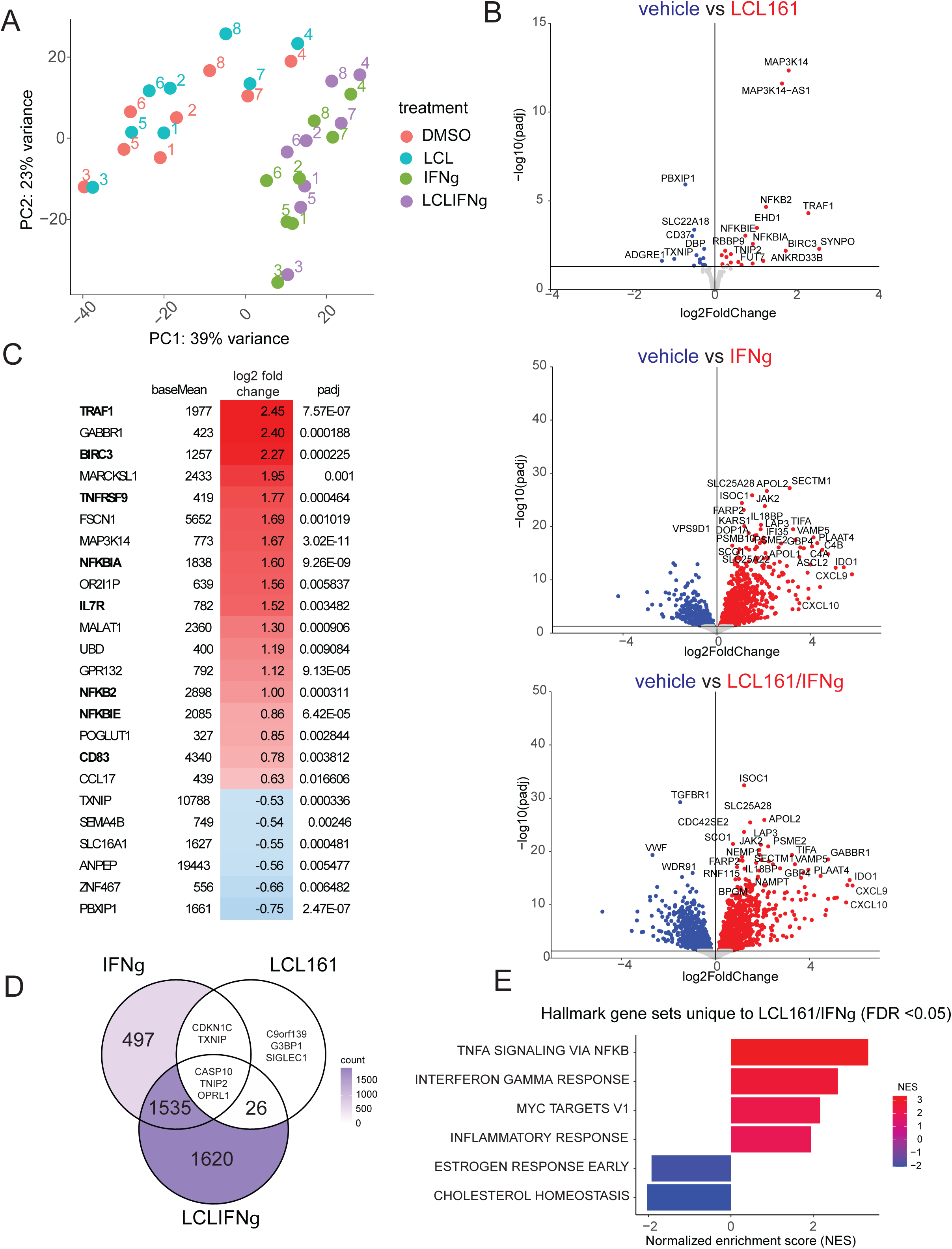
Phagocytosis is associated with a distinct transcriptional signature. A) Healthy donor macrophages from n=8 donors were treated for 24 hours with DMSO vehicle or 500nM LCL161 +/-20μg/mL IFNγ. RNA was prepared and analyzed by bulk RNAseq and principle component analysis. Individual donors are labeled with unique numbers. B) Differentially expressed genes for each of the treatment conditions compared to vehicle control. Genes with a base mean expression <50 were excluded. C) Heatmap of the top 24 most differentially expressed genes with a base mean expression >200 comparing LCL161/IFNγ versus IFNγ conditions. Base mean expression and adjusted p values are shown. D) Venn diagram showing the overlap of significant differentially expressed genes from B for each group versus vehicle controls. E) Gene set enrichment analysis for Hallmark pathways for the 1620 genes from D that were uniquely regulated in the LCL161/IFNγ treatment group.

To deconvolute the genes uniquely characterizing the phagocytic cell state, we compared combination treatment to IFNγ monotherapy. The 24 top differentially expressed genes are shown in **Figure 5C**, with members of the TNFα signaling via NF-κB Hallmark gene set listed in bold. Among this list are several upregulated genes of interest that affect macrophage cell state including CD83, a marker of activated phagocytes, and the co-stimulatory receptor 4-1BB (*TNFRSF9*).

NIK (*MAP3K14*) is not formally part of the TNFα Hallmark gene set, despite being the central regulator of the alternative NF-κB pathway ^46^. NIK is constitutively low in cells due to ubiquitination by cIAP1/2, and treatment with SMAC mimetics to degrade cIAP1 allows NIK to accumulate. NIK was not detected in our short-term proteomics analysis, this combined with the increased transcript levels indicates that new transcription is required to restore NIK levels after degradation of cIAP1 in macrophages.

IFNγ alone induces modest phagocytosis in some donors; therefore, we reanalyzed overlap of differentially expressed genes from each treated condition compared to vehicle. This method allows for genes modestly induced by IFNγ but more strongly induced by the combination therapy to be counted as unique to the combination treatment (**Figure 5D**). Gene set enrichment analysis of the resulting 1620 genes induced by combination treatment shows upregulation of NF-κB signaling, IFN response genes, and MYC targets as the dominant upregulated pathways (**Figure 5E**).

### TNFα serves as an autocrine positive feedback loop

Treatment with LCL161 leads to autocrine production of TNFα by cancer cells and by T cells ^23,29^; therefore, we reasoned that part of the strong induction of NF-κB regulated transcripts in LCL161/IFNγ treated macrophages might be due to TNFα signaling. To test this, we cultured macrophages with LCL161 and/or IFNγ in the presence of isotype control or TNFα blocking antibody. Secreted TNFα was highest in the combination treated group (**Figure 6A**). Across three different healthy donors, the degree of phagocytosis induction was reduced by TNFα blockade, suggesting that secreted TNFα contributes to a positive feedback loop to sustain the phagocytic macrophage phenotype (**Figure 6B**).

**Figure 6:**
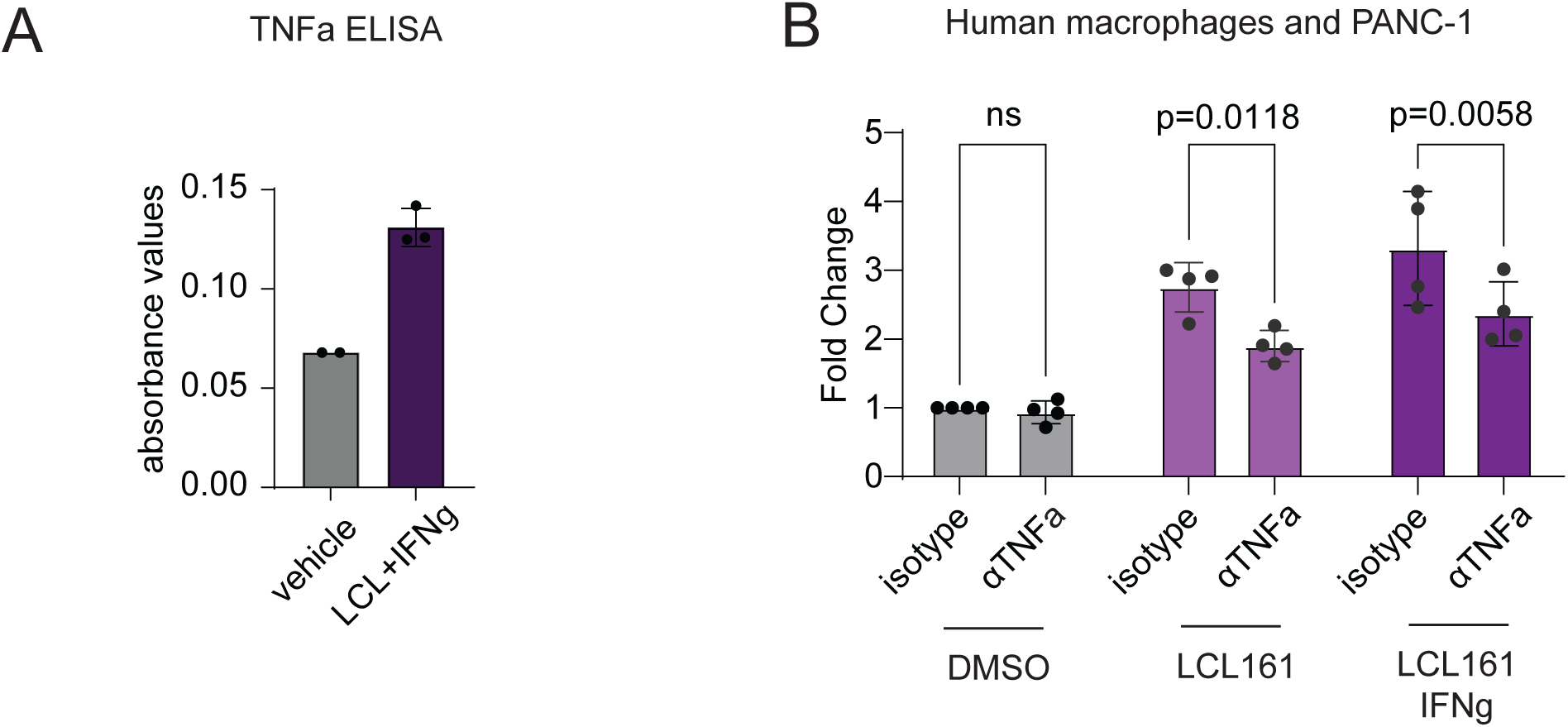
Autocrine TNFα sustains the phagocytosis phenotype. A) Healthy donor macrophages were treated with 500nM LCL161 and 20μg/mL IFNγ or vehicle control. TNFα was measured by ELISA of 24 hour culture supernatants and reported as absorbance values. Error bars are SD. B) Healthy donor macrophages from n=3 independent donors were treated with DMSO, LCL161, or LCL161+IFNγ for 18 hours of co-culture with CellTrace Violet labeled PANC-1 pancreatic cancer cells with or without TNFα blocking antibodies (1μg/mL). Phagocytosis was assessed by flow cytometry after 24 hours of co-culture. Error bars are SD. Statistical significance was determined using a pairwise one-way ANOVA with Sidak’s multiple comparisons assuming sphericity.

## Discussion

Macrophages can engulf and kill live tumor cells by phagocytosis, as shown by several groups including ours ^10,34,48,49^. We confirm and extend this finding, showing that phagocytosis is robustly induced across a range of human donors and using a range of pharmacologically distinct SMAC mimetics. Phagocytosis is further enhanced by IFNγ, a cytokine well known for its pleiotropic anti-tumor functions. These results are important to define and establish that human macrophages can be induced to phagocytosis live cancer cells. The phagocytosis phenotype involves distinct transcriptional changes and is sustained by autocrine TNFα.

Live cell phagocytosis has been challenging to demonstrate in vivo in either mice or humans. Macrophages are highly efficient phagocytes and will engulf fluorescently labeled tumor apoptotic bodies and other debris. Demonstrating that a tumor cell in vivo was alive at the time of engulfment by tumor-infiltrating macrophages is difficult with current technical capabilities, although ongoing efforts to use fluorescent reporters that are pH sensitive and activatable by caspase cleavage may be able to distinguish phagocytosis of live versus apoptotic cells in preclinical models ^50^. In the context of a human clinical trial, such fluorescent reporter systems are not possible. Functional ex vivo assays can be performed using peripheral blood monocyte-derived macrophages ^33^, although such assays do not determine whether phagocytosis is happening in the tumor itself. We therefore have heavily relied on single cell transcriptional profiling to identify phagocytic macrophages, which can be loosely defined as single cells that co-express macrophage markers and markers consistent with another cell type, such as a lymphocyte ^34^. This approach has several limitations, most notably the concern that such cells are doublets and the concern that apoptotic tumor cells may still contain RNA that can be amplified and sequenced. We therefore are highly interested in identifying a transcriptional signature of phagocytic macrophages that is not reliant on having already engulfed a live target cell. To this end, we propose a 24 gene phagocytosis signature that can be applied to single cell data sets from human cancers, although the extent to which the presence of phagocytosis-signature positive cells correlates with outcomes remains to be tested.

Therapies aimed at reprogramming macrophages have not yet demonstrated clinical success as single agents. This includes antibodies to CD47 or SIRPα, which routinely performed exceedingly well in xenograft mouse models of human cancer where the only source of human CD47 was on tumor cells and the cross-species affinity between mouse SIRPα and human CD47 is higher than the cognate interactions in either mice or humans ^51^. One lesson to be learned from the CD47 experience is that testing therapies in multiple kinds of models, including syngeneic mice and fully human ex vivo cultures, is important. Nevertheless, the discovery of CD47 as a don’t eat me signal has paved the way for other therapies aimed at inducing macrophage phagocytosis as a means of reprogramming this abundant component of the tumor microenvironment ^49^. SMAC mimetics are one approach. Other options include agonistic anti-CD40 which both activates macrophages to become tumoricidal and increases T cell activation through upregulation of costimulatory ligands on dendritic cells ^52^. We propose that combination strategies aimed at inducing both antigen-specific T cells, which provide a critical source of IFNγ, and induction of phagocytic macrophages may be optimal for sustained tumor control.

## Acknowledgements

SKD was funded by the Ludwig Center at Harvard Medical School, the Hale Family Center for Pancreatic Cancer Research at Dana-Farber, NIH R01AI158488, U01CA274276, and is a Member of the Parker Institute for Cancer Immunotherapy. SKD and MD were funded by R01AI169188. LQ was funded by the Claudia Adams Barr Foundation.

## Declaration of Interests

SKD received research funding unrelated to this project from Novartis, Bristol-Myers Squibb, Takeda, and is a founder, science advisory board member and equity holder in Kojin and has equity in Axxis Bio. MD has research funding from Eli Lilly; he has received consulting fees from Genentech, ORIC Pharmaceuticals, Partner Therapeutics, SQZ Biotech, AzurRx, Eli Lilly, Mallinckrodt Pharmaceuticals, Aditum, Foghorn Therapeutics, Palleon, and Moderna; and he is a member of the Scientific Advisory Board for Neoleukin Therapeutics, Veravas and Cerberus Therapeutics and has equity in Axxis Bio. KAD receives or has received consulting fees from Neomorph and Kronos Bio. The remaining authors declare no competing interests.

**Supplemental Figure 1:** Flow cytometry gating strategy for phagocytosis assays. A) Representative flow plots showing the phagocytosis rates gated out of total live macrophages. B) The frequency of macrophages recovered after 24 co-culture with tumor cells under each of the indicated treatment conditions.

