## Supplemental Figure 1 for "SMAC mimetics induce human macrophages to phagocytose live cancer cells"

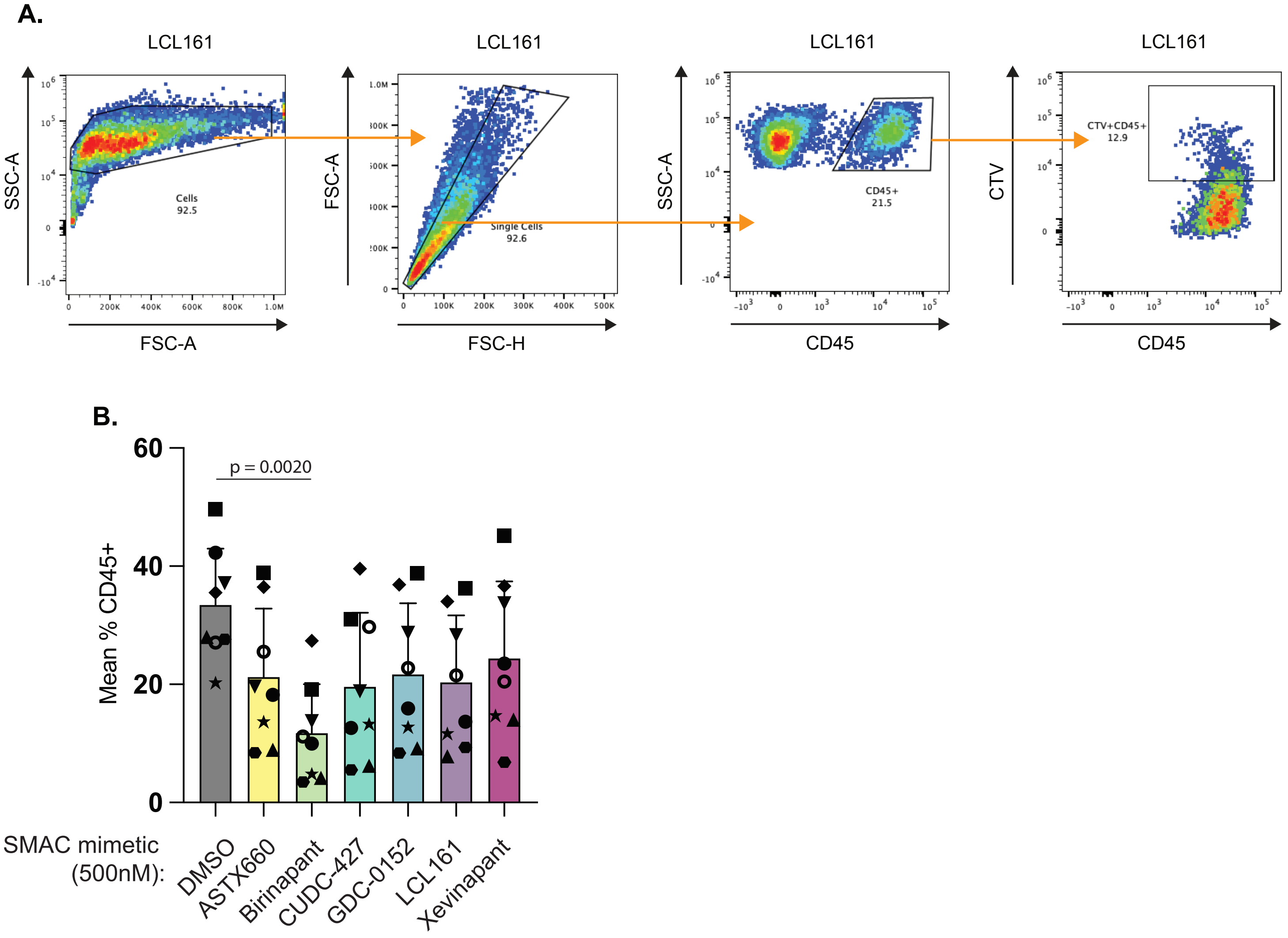


**Supplemental Figure 1: Flow cytometry gating strategy for phagocytosis assays**. A) Representative flow plots showing the phagocytosis rates gated out of total live macrophages. B) The frequency of macrophages recovered after 24 co-culture with tumor cells under each of the indicated treatment conditions.
